# *Irx1* and *Irx2* play dose-dependent cooperative functions in mammalian development

**DOI:** 10.1101/2022.10.03.510739

**Authors:** Sepideh Sheybani-Deloui, Leo Xu, Lijuan Hu, Qiongjing Yuan, Joe Eun Son, Kyoung-Han Kim, Weifan Liu, Rong Mo, Xiaoyun Zhang, Lijun Chi, Paul Delgado Olguin, Chi-Chung Hui

**Author notes:** **Present address:** Department of Cellular and Molecular Medicine, University of Ottawa, Ottawa, K1H 8M5, ON, Canada.

## Abstract

*Irx1* and *Irx2* (*Irx1/2*) are two closely linked and widely expressed members of the conserved *Iroquois* homeobox family of transcription factors. Despite mounting evidence suggesting the importance of homologs of these genes in many aspects of vertebrate development and function, the role of *Irx1/2* in mammals has remained largely unknown. Here, we used mice carrying our newly generated *Irx1*^*flox*^ and *Irx1*^*flox*^*Irx2*^*del*^ mutant alleles to perform a stepwise genetic ablation of *Irx1* and *Irx2* levels. Our analysis revealed reduced postnatal growth and viability of *Irx1*^*KO*^ mice with gross histological defects in the lung and gut and demonstrated that ablation of one copy of *Irx2* in these mice results in neonatal lethality with exacerbated phenotypic defects. Conversely, while *Irx2*^*KO*^ mice appear normal, ablation of one copy of *Irx1* in these mutants leads to lethality at weaning. Furthermore, we found that homozygous deletion of both *Irx1* and *Irx2* results in embryonic lethality by mid-gestation with defective extraembryonic vasculature. Our results illustrate that *Irx1* and *Irx2* play distinct dose-dependent cooperative functions during both the early and late stages of mouse development.

## Introduction

*Iroquois* homeobox (*Irx*) genes encode a family of homeodomain-containing transcription factors that are conserved from worms to vertebrates.^1^ In mice and humans, *Irx* genes are organized in two 3-gene clusters; *IrxA* cluster on mouse chromosome 13 (human chromosome 5) contains *Irx1, Irx2*, and *Irx4*, and *IrxB* cluster on mouse chromosome 8 (human chromosome 16) consists of *Irx3, Irx5*, and *Irx6*.^2,3^ The first two genes of each cluster display a highly overlapping expression domain while the expression of the third gene is more divergent,^4,5^ partly due to the presence of conserved shared enhancers which preferentially interact with the promoter of the cluster’s first two genes.^6^ Overexpression and RNAi or morpholino knockdown experiments in both the fly and vertebrates have illustrated the prepatterning role of *Irx* genes at early stages of development to define large territories, such as the dorsal regions of the eye, head, and mesothorax of *Drosophila*, and *Xenopus* neural ectoderm.^1,7–13^ Consistent with the expression pattern of *Irx* genes through distinct stages of development, they function again to subdivide large territories into smaller domains such as in *Drosophila* bristles, notum and wing veins, and *Xenopus* midbrain-hindbrain organization.^14–16^

Despite evidence suggesting a role of *IrxA* cluster genes in regionalization and patterning in chicks, Zebrafish and *Xenopus*, genetic studies are yet to uncover the function of *Irx1* and *Irx2* in mammals. While *Irx2* is expressed in the developing heart, nervous system, and other organs, *Irx2*-deficient mice are viable and fertile, and possess normal cardiac morphology and function.^17^ Owing to its overlapping expression with *Irx2* during embryonic development, it is probable that *Irx1* could compensate for the function of *Irx2* in *Irx2*-deficient mice. In addition, RNA-seq data from multiple stages of craniofacial/tooth development in mice from E10.5 to P4 identified *Irx1* expression at early stages of murine development.^18^ Generation of *Irx1* knockout mice (*Irx1*^-/-^) by replacing the entire *Irx1* sequence from the start codon to the stop codon by LacZ-Neo resulted in lethality shortly after birth due to defects in later stages of lung development.^18^ Thus, data from these single knockout mice studied so far have failed to reveal a role of *Irx1* and *Irx2* in embryonic development and the possibility of functional compensation has not been unaddressed.

To elucidate the mammalian functions of these evolutionarily conserved transcriptional regulators, we present here a mutational analysis of *Irx1* and *Irx2* in mice through generating compound and double knockout mice harboring cis-mutant alleles of *Irx1* and *Irx2*. We found that *Irx1*^*KO*^ mice are born with a slight reduction in their alveolar capacity and decreased intestinal villi structures, resulting in growth retardation, with majority of them dying by weaning age. Analysis of different combination of compound mutants revealed that reduction of *Irx2* dosage in the *Irx1*^*KO*^ background is sufficient to result in neonatal lethality with a greater reduction in alveolar space, which is likely not sufficient for postnatal survival. While *Irx2*^*KO*^ display no marked abnormality, reduction of *Irx1* dosage in the *Irx2*^*KO*^ background results in phenotypes similar to those of *Irx1*^*KO*^, suggesting that both copies of *Irx1* are required to compensate for the lack of *Irx2* postnatally. Furthermore, homozygous deletion of both *Irx1* and *Irx2* results in embryonic lethality by mid-gestation unveiling that at least one copy of either *Irx1* or *Irx2* is required to establish the extraembryonic vasculature that sustains growth and development before the initiation of placental blood circulation at the fetal-maternal interface. Our study illustrates a dose-dependent functional redundancy between *Irx1* and *Irx2* during the early and late stages of embryonic as well as postnatal development.

## Results

### 1. Expression of *Irx1* and *Irx2* in mice and humans

Mammalian expression of *Irx1* and *Irx2* was evaluated by integrating available data from mouse and human gene expression studies. Public scRNA-seq databases of whole mouse embryo during gastrulation from embryonic day (E) 6.5 to E8.5 revealed *Irx1* and *Irx2* expression in derivatives of all three germ layers.^19^ By analyzing the transcriptomic profile of E8.25 embryo, we found that *Irx1* and *Irx2* are detected in 80% of all the cells analyzed, predominantly in different mesodermal tissues, neuroectoderm, definitive endoderm, gut and extraembryonic endoderm, and ectoderm (**Fig. 1A**). Digital in situ platform,^20^ an image-based single-cell transcriptomics of E8.75 embryos, demonstrates the overlapping expression domains of *Irx1* and *Irx2* in the craniofacial region, cardiomyocytes, gut tube, the embryonic and extraembryonic endoderm and mesoderm (Supplemental Fig.). Furthermore, data from the Mouse Genome Informatics (MGI)^21^ database suggests that *Irx1* and *Irx2* continue to be expressed in organs required for growth and survival such as the gut, heart, lung, and kidney during organogenesis and postnatally (**Fig. 1B**). We also took advantage of bulk RNA-seq and scRNA-seq databases from the human protein atlas (HPA)^22,23^ to confirm that the expression domains of murine *Irx1* and *Irx2* are conserved in human organs, with the exception of intestine (Fig. 1C, D). The conserved overlapping expression domain of *Irx1* and *Irx2* prompted us to investigate their cooperative function by stepwise genetic ablation of these two genes.

**Figure 1.**
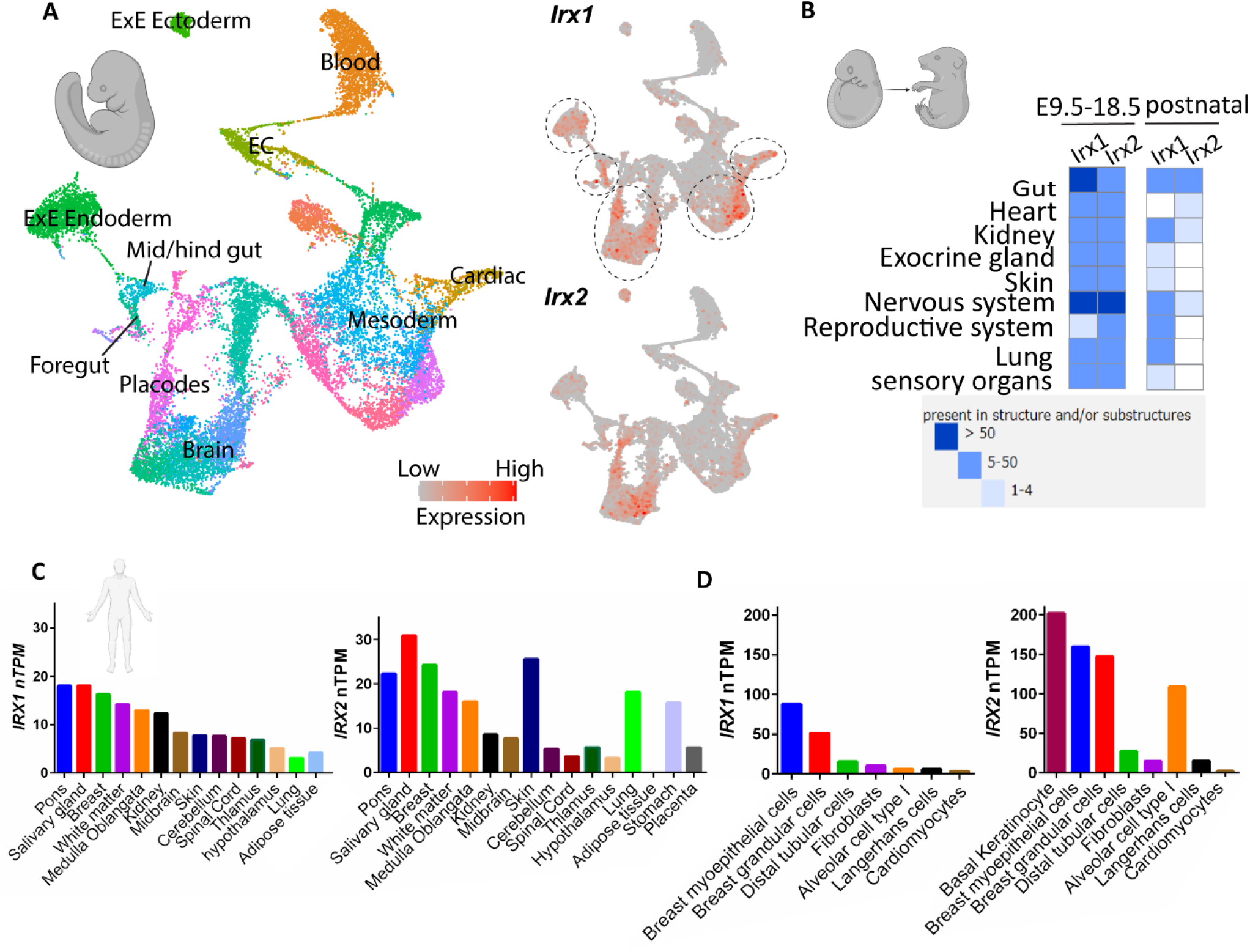
Overlapping expression of *Irx1* and *Irx2* in derivatives of all three germ layers. **A**. Uniform manifold approximation and projection (UMAP) plot showing clusters of cells from E8.25 embryo single-cell transcriptomic atlas data set generated by Marioni lab at the Cancer Research UK Cambridge Institute. Cells are colored by their annotated cell type, and groups of major embryonic and extraembryonic lineages are annotated on the UMAP. UMAP plots showing *Irx1* and *Irx2* expressing cells, with color intensity proportional to log2 normalized counts for each transcript. **B**. Matrix of *Irx1* and *Irx2* expression across different organs in post-gastrulation embryo until birth, as well as postnatal period based on Mouse Genome Informatics database. The blue color in the grid denotes the presence of evidence for expression and gets progressively darker when there is more supporting evidence. **C**. Expression of *IRX1* and *IRX2* across human tissue based on Human Protein Atlas collection of human protein atlas (HPA) bulk tissue RNA-seq data reported as normalized transcript per million (nTPM). **D**. Expression of *IRX1* and *IRX2* in single cell types based on HPA collection of single-cell RNA sequencing (scRNAseq) data from 25 human tissues and peripheral blood mononuclear cells reported as nTPM. Data retrieved from www.proteinatlas.org.

### 2. Generation of *Irx1* and *Irx2* single-knockout mouse models

Because the distance between the *Irx1* and *Irx2* loci is around 0.6 Mbps, we performed sequential gene targeting to study the function of *Irx1* and *Irx2 in vivo*. We first generated a conditional *Irx1* allele by CRISPR/Cas9-mediated insertion of *lox*P sequences to flank the second exon of *Irx1* without affecting its reading frame. The resulting *Irx1*^*flox*/+^ mouse line was crossed with a mouseline expressing a ubiquitous CRE-recombinase, resulting in *Irx1*^*del*/+^ mice, which were then intercrossed to generate *Irx1*^*del*/*del*^ (called *Irx1*^*KO*^, hereafter) (**Fig. 2A**). Both *Irx1*^*flox*/+^ and *Irx1*^*del*/+^ mice appear normal and show normal fertility. We confirmed the insertion of the *lox*P site and deletion of the *lox*P-flanked exon in *Irx1*^*del*/+^ mice, respectively, using PCR with primers designed specifically to discern these alleles (**Fig. 2B**). qRT-PCR analysis using primers specific to the site of deletion confirms largely diminished levels of the exon 2 transcript of *Irx1* in *Irx1*^*KO*^ embryos, demonstrating the generation of a null allele (**Fig. 2C**).

**Figure 2.**
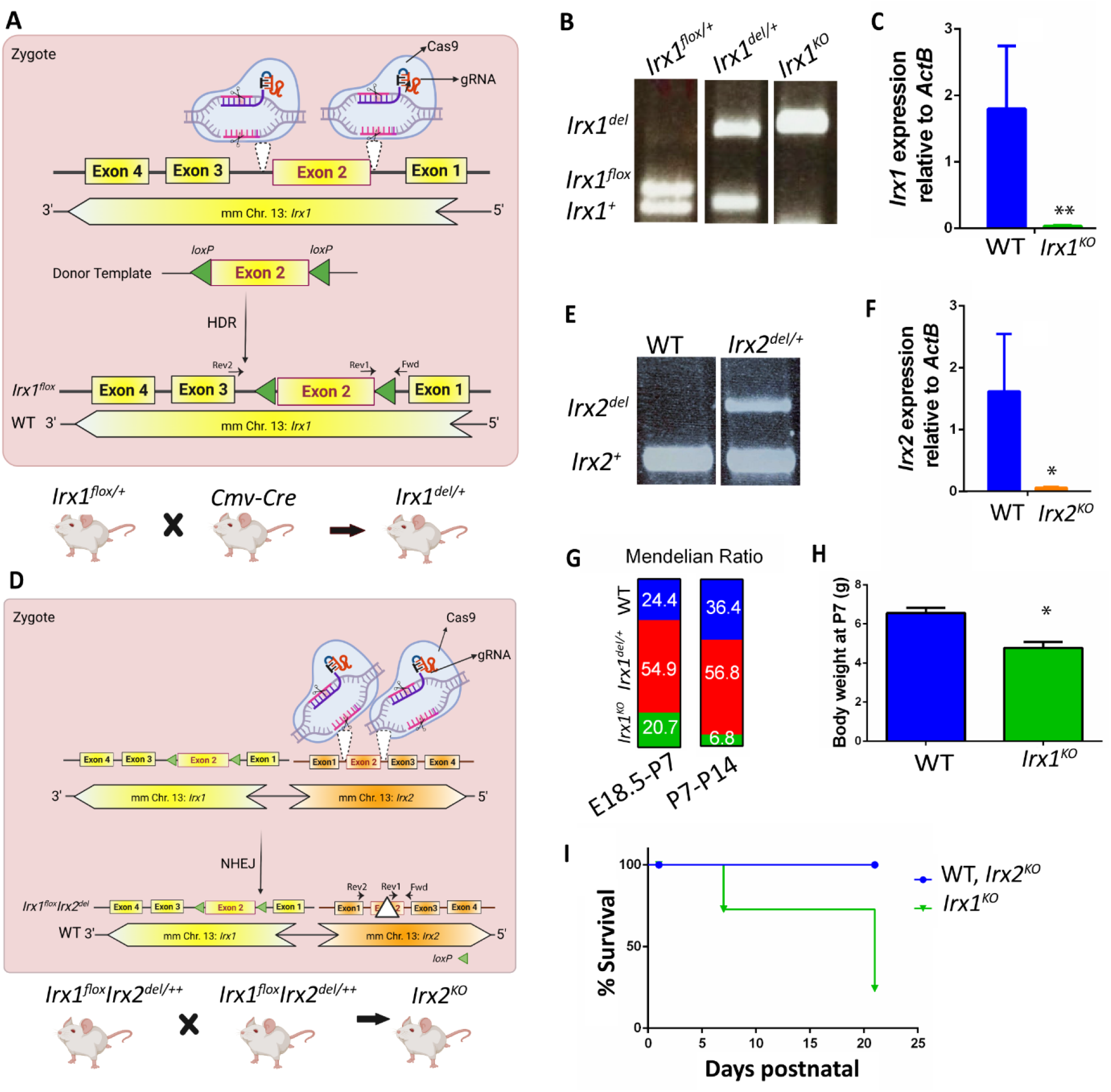
*Irx1* is required for postnatal mouse survival while *Irx2* is dispensable. **A**. Schematic depicting the *IrxA* locus and CRISPR-guided gene insertion in the zygote to generate *lox*P flanked *Irx1*. Breeding strategy to generate mice with whole body homozygous deletion of *Irx1* using a ubiquitous Cre. Arrows mark the location of primers to detect the wild type, *lox*P flanked, and mutant *Irx1* alleles. HDR: homologous direct repair **B**. PCR result showing successful deletion of exon 2 after Cre-*lox*P recombination. **C**. qRT-PCR analysis of *Irx1* mRNA level relative to ActinB in WT and *Irx1*^*KO*^ whole embryos at E9.5. **D**. Schematic depicting the *IrxA* locus and CRISPR-guided deletion to generate *Irx2* mutant allele on the chromosome harboring *lox*P flanked *Irx1*. Breeding strategy to generate homozygous deletion of *Irx2*. Arrows mark the location of primers to detect wildtype and mutant *Irx2* alleles. NHEJ: non-homologous ends joining. **E**. PCR result showing successful deletion of *Irx2*-exon2 in CRISPR-CAS9 products. **F**. qRT-PCR analysis of *Irx2* mRNA level relative to ActinB in WT and *Irx2*^*KO*^ whole embryos at E9.5. **G**. Mendelian ratio of each genotype among pups recovered between E18.5 and P7 from intercrossing *Irx1*^*del*/+^ mice, compared to the ratio among those recovered between P7-P14. **H**. Average body weight of *Irx1*^*KO*^ compared to WT control pups at P7. **I**. Survival curve over the postnatal period comparing WT, *Irx2*^*KO*^, and *Irx1*^*KO*^ survival rates.

We next used zygotes that emerged from *Irx1*^*flox*/*flox*^ intercrosses to introduce a CRISPR/Cas9-mediated deletion of the second exon of *Irx2* on the same chromosome harboring *Irx1*^*flox*^, resulting in *Irx1*^*flox*^*Irx2*^*del*^/+ mice (**Fig. 2D**). By intercrossing *Irx1*^*flox*^*Irx2*^*del*^/+ mice, we generated *Irx2*^*KO*^ (*Irx1*^*flox*^*Irx2*^*del*^/*Irx1*^*flox*^*Irx2*^*del*^) mice and confirmed their genotype using PCR to amplify the *Irx1*^*flox*^ and *Irx2*^*del*^ alleles (**Fig. 2E**). In addition, qRT-PCR analysis using primers specific to the site of deletion confirms lack of the exon 2 transcript of *Irx2* in *Irx2*^*KO*^ embryos (**Fig. 2F**), demonstrating the successful generation of an *Irx2* deletion allele in *cis* with the floxed allele of *Irx1*.

Consistent with our previously published report, *Irx2*^*KO*^ mice are viable and fertile and show no abnormality. However, contrary to the previous report of perinatal lethality in *Irx1*-deficient mice,^18^ we observed a reduction in the percentage of surviving *Irx1*^*KO*^ pups only after Postnatal day (P) 7 (**Fig. 2G**). *Irx1*^*KO*^ mice are significantly smaller than WT littermate but they are still recovered at a 20% Mendelian ratio at P7 (**Fig. 2H**). Survival curve comparison shows a 75% reduced viability of *Irx1*^*KO*^ mice compared to *Irx2*^*KO*^ and WT mice by weaning age (**Fig. 2I**).

### 3. Dose-dependent functional redundancy between *Irx1* and *Irx2*

We conducted a histological analysis of vital organs with endogenous *Irx1*/2 expression, including the lung, intestine, and heart, soon after birth. Our data shows no marked abnormality in *Irx2*^*KO*^ lungs while measurement of alveolar airspace using ImageJ shows a 35% reduction in airspace in *Irx1*^*KO*^ mice (*p*-value<0.01) (**Fig. 3A**). Similarly, while there is no visible phenotype in *Irx2*^*KO*^ intestines, *Irx1*^*KO*^ intestines exhibit shortened villi and mucosal shedding resulting in a relatively flat intestinal mucosa surface (**Fig. 3B**). In addition, we found a subtle thinning of the compact layer of the heart ventricular wall in *Irx1*^*KO*^ mice (**Fig. 3C**). Together, our data are consistent with the report by Lebel *et al*. (2003), suggesting that, in the presence of *Irx1, Irx2* is dispensable for development, and while *Irx1*-deficiency does not result in neonatal lethality, *Irx1* is required for the formation of a fully functional lung and intestine.

**Figure 3.**
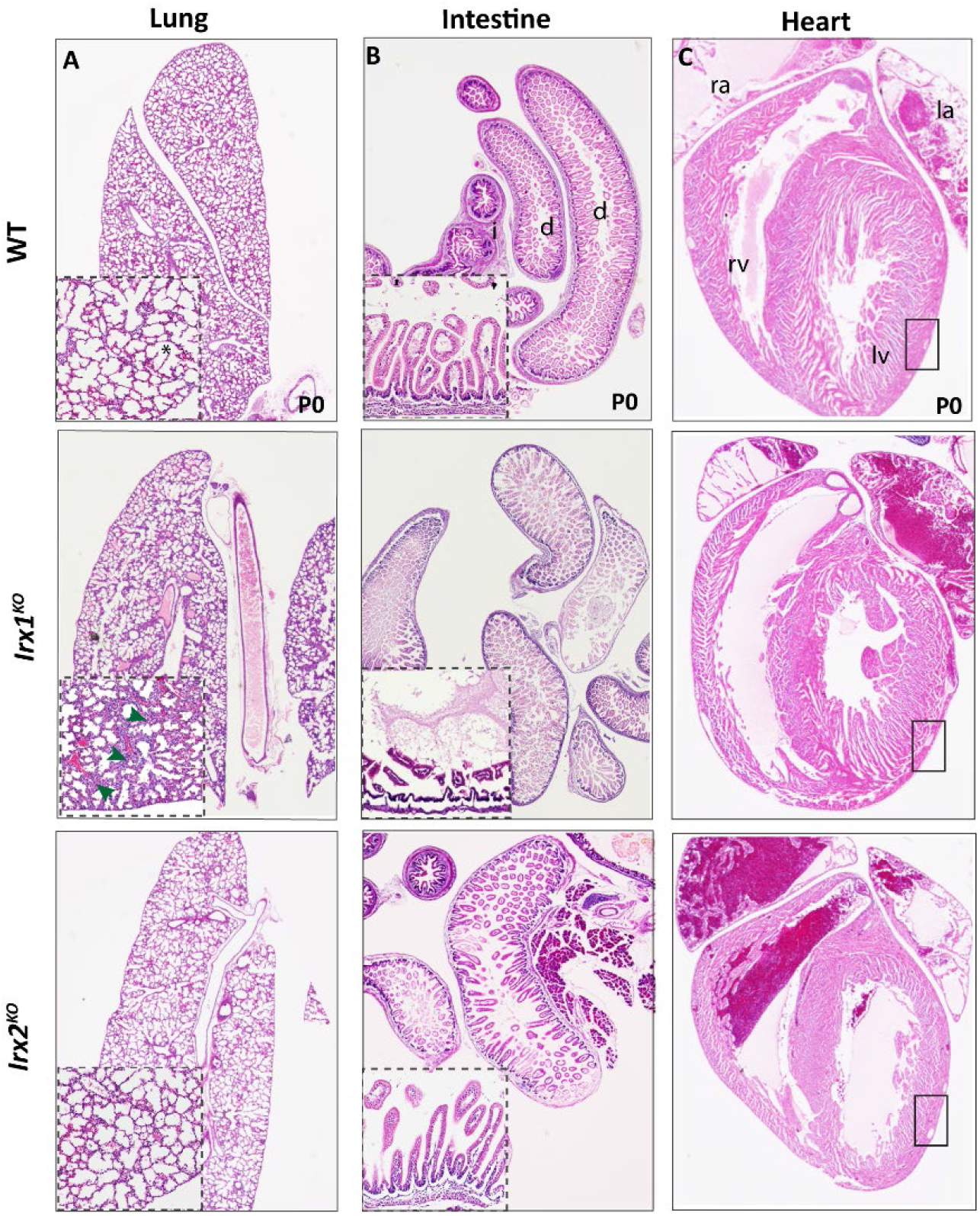
*Irx1* is required for alveolar inflation and intestinal villi maintenance. **A**. Representative images of hematoxylin & eosin (H&E) staining of the lung in WT, *Irx1*^*KO*^, and *Irx2*^*KO*^ mice shortly after birth at P0. Insets contain magnified areas of interest. Asterisk marks an inflated alveolar sac. Arrowheads point to the increased interstitial space between smaller alveolar sacs in *Irx1*^*KO*^ lungs. **B**. Representative images of H&E staining of the intestinal mucosa. Arrowheads point to the epithelial shedding and shortened intestinal villi in *Irx1*^*KO*^ mice. d:duodenum; i:ileum. Insets contain 5 times magnified areas of interest. **C**. Representative images of H&E staining of heart in WT, *Irx1*^*KO*^, and *Irx2*^*KO*^ mice at P0. Square boxes outline the thinning of the right ventricular wall in *Irx1*^*KO*^ mice. rv: right ventricle, ra: right atrium, lv: left ventricle, la: left atrium.

To determine if the removal of one copy of *Irx1* in *Irx2*-null background, or one copy of *Irx2* in *Irx1*-null background exacerbates the phenotypes described above, we crossed *Irx1*/2^*dhet*^ mice harboring the cis-heterozygous mutant alleles of *Irx1* and *Irx2* with either *Irx2*^*KO*^ or *Irx1*^*het*^ mice to generate *Irx1*^*het*^*Irx2*^*KO*^, or *Irx1*^*KO*^*Irx2*^*het*^ mice respectively (**Fig. 4A**). Intriguingly, we found that *Irx1*^*KO*^*Irx2*^*het*^ mice die shortly after birth (**Fig. 4B, C**). We confirmed that pups were initially born alive as their lung was inflated with air and floated on the water. In addition, while *Irx1*^*het*^*Irx2*^*KO*^ mice show only a 50% reduction in viability by weaning age, mice that survive are significantly smaller than their littermate controls (**Fig. 4B, D**). Histological comparison shows a further reduction in the alveolar space and a corresponding increase in the interstitial space upon removal of a single dose of *Irx2* in the *Irx1*^*KO*^ background, resulting in a 70% reduction in alveolar space (*p*-value<0.0001, **Fig. 4E**). Furthermore, *Irx1*^*het*^*Irx2*^*KO*^ mice exhibit similar histological features found in *Irx1*^*KO*^ mice in terms of intestinal villi sloughing and thinning of the sub-mucosal and muscular layers. Importantly, *Irx1*^*KO*^*Irx2*^*het*^ mice have a more severe intestinal phenotype with a relatively unfolded mucosa and a further reduction in submucosal and muscle thickness (**Fig. 4F**). Similarly, *Irx1*^*KO*^*Irx2*^*het*^ hearts exhibit further disorganization and loosening of the compact ventricular myocardium (**Fig. 4G**). These findings demonstrate a dose-dependent functional redundancy between *Irx1* and *Irx2* such that one copy of either *Irx1* or *Irx2* seems to be sufficient for embryonic function whereas both copies of at least one of the genes are required for postnatal maturation and function.

**Figure 4.**
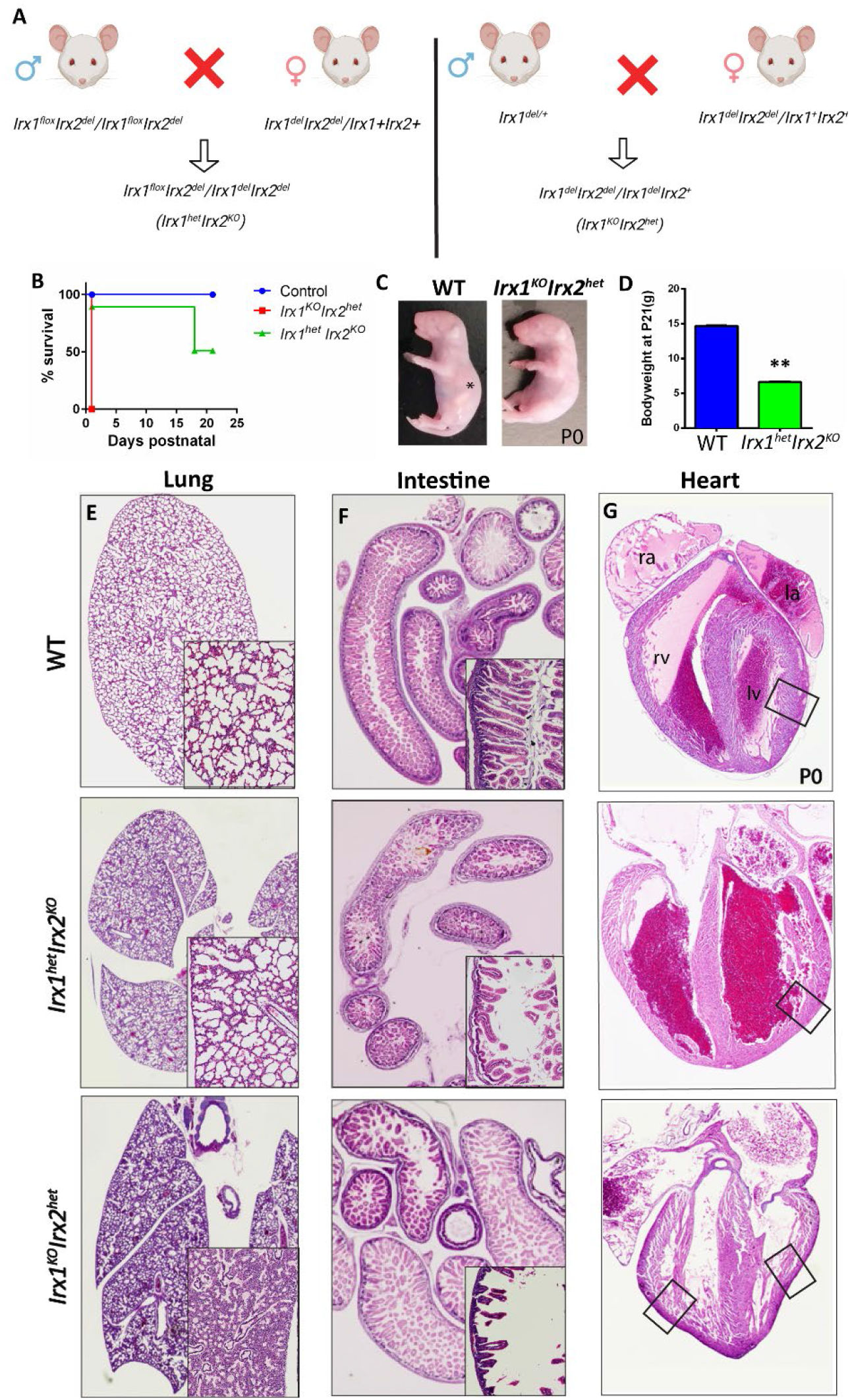
In the absence of *Irx1*, both copies of *Irx2* are required for postnatal survival. **A**. Breeding strategy to generate mice that are heterozygous for *Irx2* in the *Irx1*^*KO*^ background, and those that are heterozygous for *Irx1* in the *Irx2*^*KO*^ background. **B**. Survival curve of *Irx1*^*KO*^*Irx2*^*het*^, *Irx1*^*het*^*Irx2*^*KO*^, and WT control mice. **C**. Body weight of WT and *Irx1*^*het*^*Irx2*^*KO*^ mice that reach weaning age. **D**. Representative image of *Irx1*^*KO*^*Irx2*^*het*^ and control mice right after birth. Asterisk marks the milk-containing stomach in the control pup. **E**. Representative images of hematoxylin & eosin (H&E) staining of the lung. **G**. Representative images of H&E staining of the intestinal mucosa. Insets contain 5 times magnified areas of interest. **H**. Representative images of H&E staining of heart. Square boxes outline the thinning of the right ventricular wall.

### 4. *Irx1* and *Irx2* play a vital role in embryonic growth and survival

We next intercrossed *Irx1*^*del*^*Irx2*^*del*^/*Irx1*^+^*Irx2*^+^ mice to generate mutants with homozygous deletion of both *Irx1* and *Irx2*, called *Irx1*/2^*DKO*^ hereafter (**Fig. 5A**). Using antibodies specific to *Irx1* and *Irx2*, we demonstrated their expression in both the extraembryonic mesoderm and the underlying endoderm layer of the yolk sac, while the nucleated fetal erythroid cells showed no expression of *Irx1* or *Irx2* (**Fig. 5B**). These immunostainings were mostly absent in *Irx1*/2^*DKO*^ tissues confirming the specificity of the antibody. Moreover, we confirmed the absence of *Irx1* and *Irx2* expression in *Irx1*/2^*DKO*^ using western blot analysis on E9.5 embryos (**Fig. 5C**). We found that *Irx1*/2^*DKO*^ embryos were fully resorbed by E15.5 and no viable embryo can be recovered after E12.5 (**Fig. 5D**). Notably, a comparison of embryo size and developmental milestones such as neural closure, embryo turning and initiation of heartbeat shows no marked difference between the WT and mutants at E9.5 and E10.5, while the embryos appear pale and developmentally arrested beginning from E11.5 (**Fig. 5E**). Examining the extraembryonic structures, required for embryonic survival and growth until mid-gestation, revealed a visible lack of vasculature, including the major vitelline artery along the yolk sac which is developed through the process of vascular remodeling to provide nutrient and gas exchange to and from the embryo. These observations indicate that *Irx1* and *Irx2* share an overlapping function during yolk sac angiogenesis to support embryonic survival.

**Figure 5.**
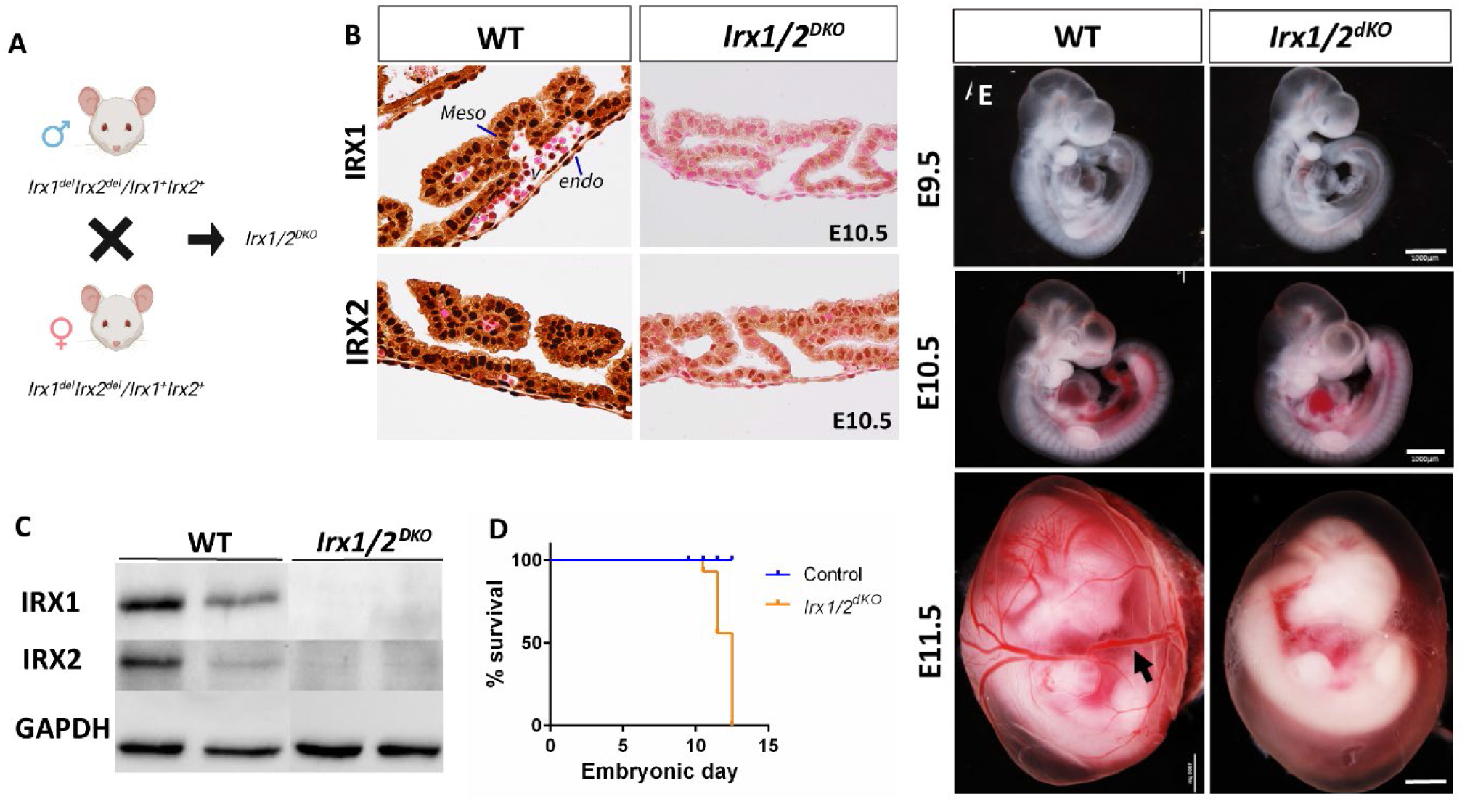
At least one copy of either *Irx1* or *Irx2* is required for embryonic growth and survival past mid-gestation. **A**. Breeding strategy to generate homozygous deletion of both *Irx1* and *Irx2*. **B**. Immunohistochemistry of yolk sac tissue isolated from WT and *Irx1*/2^*DKO*^ embryo at E10.5 using antibodies against *Irx1* and *Irx2*. Nuclei are counterstained with Eosin in red. Meso:mesoderm; Endo:endoderm; V:vessel **C**. Western blot analysis of *Irx1* and *Irx2* levels in two *Irx1*/2^*DKO*^ and two WT whole embryos at E9.5. **D**. survival curve of *Irx1*/2^*DKO*^ and control embryos during embryonic stages. **E**. Representative images of whole WT and *Irx1*/2^*DKO*^ embryos recovered at E9.5, E10.5, and E11.5. Arrowhead points to the vitelline artery present in the yolk sac of the WT embryo which is absent from the mutant embryo (scale bars = 1000μm).

## Discussion

Genetic compensation by activation of one or more closely related genes is a well-established mechanism that is thought to have evolved to ensure proper development even if one or more genes fail to function. In this study, CRISPR/Cas9-mediated stepwise ablations of *Irx1* and *Irx2* have revealed a dose-dependent genetic compensation between the first two members of the *IrxA* cluster in key organs required for mouse development, growth, and survival. Based on our analysis of public expression databases, it is evident that *Irx1* and *Irx2* are both expressed in the murine embryonic and postnatal lung, gut, and heart. In addition, *Irx1* and *Irx2* transcripts are detected in cardiomyocytes and alveolar type I cells in the human scRNAseq datasets. The broad expression of *Irx1* and *Irx2* in the foregut and midgut endoderm (the common developmental origin of both the digestive and respiratory systems), as well as the involvement of intricate molecular cross-talks between the endoderm and the surrounding mesoderm, do not allow us to identify the precise function of *Irx1* and *Irx2* in each of these lineages. In future studies, this can be addressed by crossing our *Irx1*^loxP^ mice with lineage-specific Cre or tamoxifen-inducible Cre lines to delineate the cooperative functions of *Irx1*/2 in a temporal and spatially specific manner.

In this study, we showed that *Irx1* is essential for intestinal villus development and/or maintenance, such that *Irx1*^*KO*^ mice show severe postnatal growth retardation and lower viability by weaning age. Our analysis of *Irx1*^*KO*^*Irx2*^*het*^ mice revealed that, in the absence of *Irx1*, both copies of *Irx2* are required for the respiratory and digestive systems of the mutants to sustain survival until weaning age. Although the lung dissected from *Irx1*^*KO*^*Irx2*^*het*^ pups are inflated with air, the alveolar airspace is largely reduced at the expense of increased interstitial space, and there is more severe retardation of intestinal villus formation than in *Irx1*^*KO*^ mice. During the final stage of embryonic and early postnatal development of the lung, the interstitium between the air spaces becomes thinner as the result of decreased collagen fiber deposition, while elastic fibers are deposited in the interstitium.^24,25^ This process is thought to lay the foundation for subsequent secondary septation and the formation of mature alveoli. In contrast to the extensive knowledge of the regulatory mechanisms of branching morphogenesis, it has been challenging to identify the molecular mechanisms that regulate this final stage of alveologenesis partly because majority of mouse mutants suffer from the cessation of lung development and/or death before the initiation of sacculation and alveolarization.^26^ However, in our mutants, the earlier stages of lung development are not negatively affected as we do not observe any marked histological difference in the E15.5 lung in our single and compound mutants, at the conclusion of the pseudoglandular stage of lung development. Investigation of ECM deposition, cell proliferation, and apoptosis in the interstitial space of the compound mutant mice prior to birth will reveal whether *Irx1*/2 is involved in the regulation of signaling mechanisms in sacculation and alveolarization. Similarly, between E15.5 and birth in gut, hedgehog signaling promotes a proximal to distal progression of cytodifferentiation to convert the initially pseudostratified epithelium to a monolayer of the columnar epithelium as villi are formed that are separated from each other by proliferating intervillous epithelium.^27–29^ Our data suggest that *Irx1* and *Irx2* may be cooperatively involved in the proper emergence or maintenance of villi. To examine this possibility, targeted deletion of *Irx1* followed by studying the proliferation and molecular specification of crypt and villus cells prior to birth is essential.

The difference between the phenotype of our *Irx1*^*KO*^ mice and that of a previous publication indicating neonatal lethality in the absence of *Irx1* is likely due to the difference in the molecular nature of the mutations. In the previous mutant, the entire coding sequence of *Irx1* was replaced by *lacZ*, resulting in no *Irx1* mRNA being transcribed.^18^ However, our mutants lack only the second exon of *Irx1*. It is possible that this will result in non-sense mediated mRNA decay, whereby RNA degradation intermediates could trigger genetic compensation by the closely-related *Irx2* gene. Through analysis of several models of transcriptional adaptation in zebrafish and mice, it is known that alleles that fail to transcribe the mutated gene do not display transcriptional adaptation and exhibit more severe phenotypes.^30,31^ Our hypothesis is further supported by the observation that *Irx1*^*KO*^*Irx2*^*het*^ mice display a phenotype more similar to that of the previously published *Irx1*^*KO*^ mouse model.

Although our analysis demonstrated that *Irx1*/2^*DKO*^ yolk sacs can initiate vasculogenesis and hematopoiesis, we found that both processes are deficient in their ability to undergo vascular remodeling. At E10.5 to E11.5, *Irx1*/2^*DKO*^ yolk sacs are paler than those of heterozygous and wild-type littermates, most probably owing to reduced circulation of erythrocytes in mutants. Yolk sac endothelial cells and erythrocytes have a close developmental origin as they concomitantly arise from a bipotent mesodermal population of cells called hematoendothelial progenitor cells, which receives differentiation and proliferation cues from the underlying extraembryonic mesoderm and visceral endoderm.^32–34^ Results from spatial single-cell transcriptomic atlas of mouse embryos suggest that *Irx1*/2 are expressed in hematoendothelial progenitors at E8.75 before the onset of angiogenesis defects in *Irx1*/2^*DKO*^ mutants.^20^ It will be intriguing to study how *Irx1*/2 regulate endothelial versus erythrocyte differentiation to ultimately establish vascular integrity and organization.

In summary, we generated new genetic tools to study the compensatory and unique functions of *Irx1* and *Irx2*, two poorly understood members of the *Irx* family of transcription factors. Our study demonstrates a cooperative role for *Irx1* and *Irx2* in the late lung, gut, and heart development and postnatal function. Reduced dosage of *Irx2* in the absence of *Irx1* exacerbates the alveolar airspace, gut villus, and ventricular wall deficiencies observed in *Irx1*^*KO*^ mice. Our results uncover a new developmental role for *Irx1* and *Irx2* in establishing extraembryonic circulation which is necessary for embryonic growth and survival during mid-gestation. This study forms the foundation for future studies to decipher the lineage- and temporal-specific molecular actions of *Irx1* and *Irx2* that underlies the observed phenotypes.

## Materials and Methods

### Animals

All animal experimental protocols approved by the Animal Care Committee of The Centre for Phenogenomics conformed to the standards of the Canadian Council on Animal Care. The colony was housed in a specific pathogen-free (SPF) facility in 96 ventilated cages with controlled environment settings (21-22 ºC, 30%-60% humidity) 12-h light/dark cycles, and free access to water. Mouse model generation including design and synthesis of reagents for genome editing, microinjection, and screening of the founders for the mutation of interest was performed by model production services at TCP. Founder mice were then bred with outbred CD1 line for multiple generations before selection for experimental procedures. For timed breeding, the noon of the day of a vaginal plug was considered as E0.5.

### Histology

Paraffin-embedded tissues were prefixed in 4% paraformaldehyde overnight at 4°C and sectioned at 5 μm thickness. Sections were deparaffinized and rehydrated and were either stained with hematoxylin and eosin (H&E) or boiled with sodium citrate for antigen retrieval. Sections were then blocked with donkey serum, incubated with Irx1 (Sigma, 1:100) or Irx2 antibodies^17^ (1/50) overnight at 4°C, and then with an anti-rabbit secondary antibody coupled to biotin. We used the standard Vectastain ABC-AP kit (Vector) and the red substrate kit (Vector) to visualize the signal. Images were captured using a Nikon eclipse microscope and analyzed using ImageJ (National Institutes of Health (NIH).

### Gene expression analysis

Total RNA was extracted from embryos using RNeasy Mini (Qiagen) and complementary DNA was synthesized using M-MLV reverse transcriptase (Thermofisher) with oligo(dT). Gene expression assay was conducted using SYBR Green methods on Applied Biosystems 7900 HT fast RT-PCR system, and relative cycle threshold (ddCT) values were normalized by *Actb* (β-actin). Primers for specific transcripts were designed using RNA blast and verified through UCSC in silico PCR.

### Single-cell RNA-seq data analysis

The processed and annotated mouse gastrulating embryo scRNA-seq dataset was downloaded from Array Expression, E-MTAB-6967. We used Seurat v3 pipelines to identify significant PC dimensionalities by the ElbowPlot method and used the ‘RunUMAP()’ function to conduct UMAP analysis to reduce the significant PCs to two dimensions. Expression of genes of interest was then visualized using the ‘FeaturePlot’ function.

---

**Supplemental Figure.**
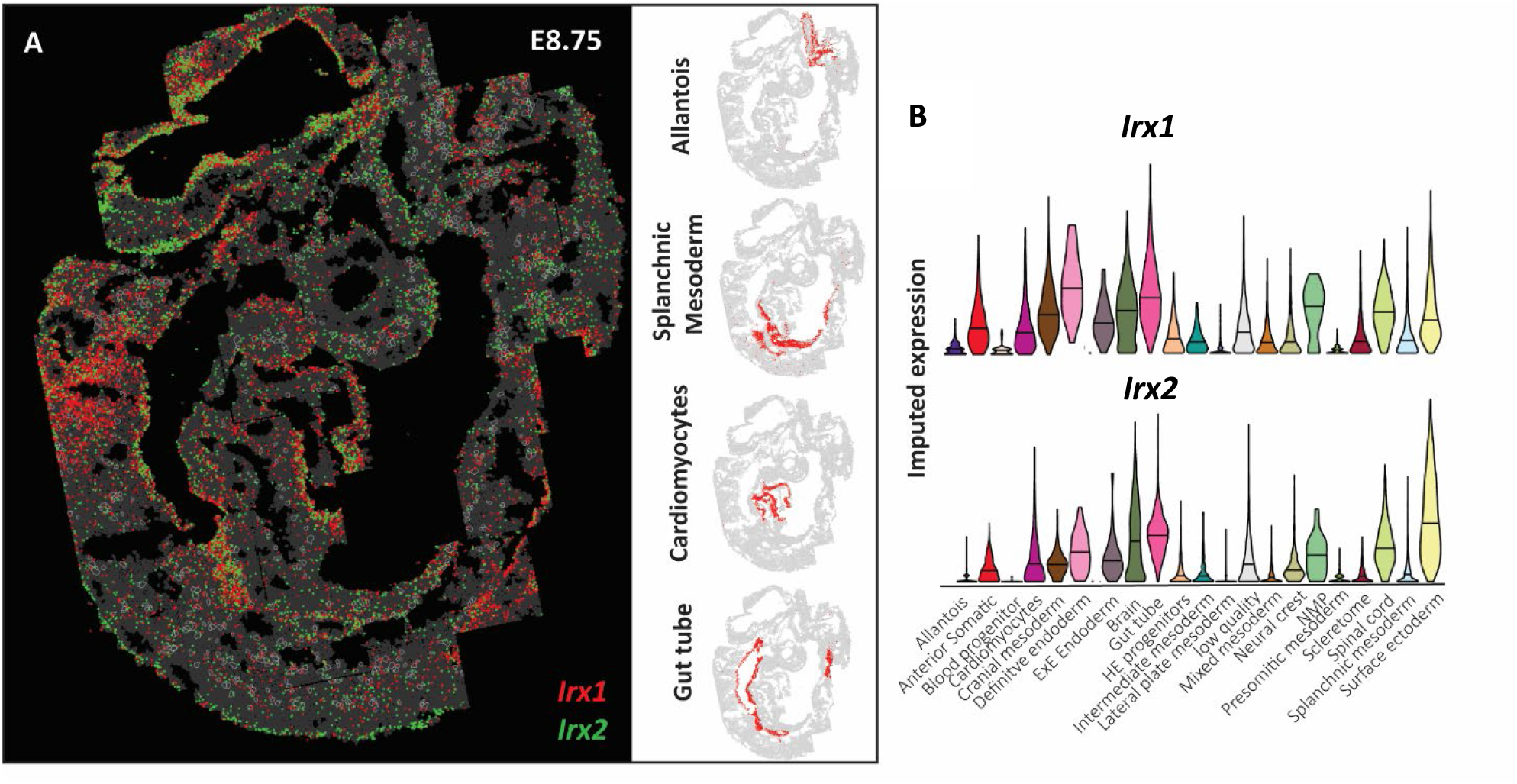
Spatial characterization of *Irx1/2* expression during mouse organogenesis. **A**. Digital *in situ* showing detected mRNA molecules for Irx1 (red) and Irx2 (green) across the entire E8.75 embryo sections. Areas pertaining to the allantois, splanchnic mesoderm, cardiomyocytes, and gut tube are highlighted in red on the right..B. Imputed gene expression of *Irx1* and *Irx2* across different cell types based on the integration of multiplexed transcriptional measurements with two single-cell transcriptome atlases (Data available from https://crukci.shinyapps.io/SpatialMouseAtlas/)

